# Effect size and statistical power in the rodent fear conditioning literature – a systematic review

**DOI:** 10.1101/116202

**Authors:** Clarissa F. D. Carneiro, Thiago C. Moulin, Malcolm R. Macleod, Olavo B. Amaral

## Abstract

Proposals to increase research reproducibility frequently call for focusing on effect sizes instead of p values, as well as for increasing the statistical power of experiments. However, it is unclear to what extent these two concepts are indeed taken into account in basic biomedical science. To study this in a real-case scenario, we performed a systematic review of effect sizes and statistical power in studies on learning of rodent fear conditioning, a widely used behavioral task to evaluate memory. Our search criteria yielded 410 experiments comparing control and treated groups in 122 articles. Interventions had a mean effect size of 29.5%, and amnesia caused by memory-impairing interventions was nearly always partial. Mean statistical power to detect the average effect size observed in well-powered experiments with significant differences (37.2%) was 65%, and was lower among studies with non-significant results. Only one article reported a sample size calculation, and our estimated sample size to achieve 80% power considering typical effect sizes and variances (15 animals per group) was reached in only 12.2% of experiments. Actual effect sizes correlated with effect size inferences made by readers on the basis of textual descriptions of results only when findings were non-significant, and neither effect size nor power correlated with study quality indicators, number of citations or impact factor of the publishing journal. In summary, effect sizes and statistical power have a wide distribution in the rodent fear conditioning literature, but do not seem to have a large influence on how results are described or cited. Failure to take these concepts into consideration might limit attempts to improve reproducibility in this field of science.

## Introduction

Biomedical research over the last decades has relied heavily on the concept of statistical significance – i.e. the probability that an effect equal to or larger than that observed experimentally would occur by chance under the null hypothesis – and classifying results as “significant” or “non-significant” on the basis of an arbitrary threshold (usually set at p < 0.05) has become standard practice in most fields. This approach, however, has well-described limitations that can lead to erroneous conclusions when researchers rely on p values alone to judge results [1–6]. First of all, p values do not measure the magnitude of an effect, and thus cannot be used by themselves to evaluate its biological significance [7]. Moreover, the predictive value of a significance test is heavily influenced by factors such as the prior probability of the tested hypothesis, the number of tests performed and their statistical power [8]; thus, similar p values can lead to very different conclusions in distinct scenarios [1].

Recent calls for improving research reproducibility have focused on reporting effect sizes and confidence intervals alongside or instead of p values [6–9] and for the use of both informal Bayesian inference [10] and formal data synthesis methods [11] when aggregating data from multiple studies. The concepts of effect size and statistical power are central for such approaches, as how much a given experiment will change a conclusion or an effect estimate will depend on both. However, it is unclear whether they receive much attention from authors in basic science publications. Discussion of effect sizes seems to be scarce, and recent data has shown that sample size and power calculations are very rare in the preclinical literature [12,13]. The potential impact of these omissions is large, as reliance on the results of significance tests without consideration of statistical power can decrease the reliability of study conclusions [14].

Another issue is that, if effect size is not taken into account, it is difficult to adequately assess the biological significance of a given finding. As p values will be low even for small effect sizes if sample size is large, biologically trivial effects can be found to be statistically significant. In preclinical studies, overlooking effect sizes will thus lead to inadequate assessment of therapeutic potential, whereas in basic research it will cause difficulties in dissecting essential biological mechanisms from peripheral modulatory influences [15]. The wealth of findings in the literature will thus translate poorly into better comprehension of phenomena, and the abundance of statistically significant findings with small effect sizes can eventually do more harm than good. This problem is made much worse when many of these studies have low positive predictive values due to insufficient power, leading a large fraction of them to be false positives [8,14,16–18].

To analyze how effect sizes and statistical power are taken into account in the description and publication of findings in a real-case scenario of basic biomedical science, we chose to perform a systematic review of articles on learning of rodent fear conditioning, probably the most widely used behavioral task to study memory in animals [19]. Focusing on this task provides a major advantage in the fact that the vast majority of articles use the same measure to describe results (i.e. percentage of time spent in freezing behavior during a test session). As effect sizes are comparable across studies, studying their distribution allows one to estimate the statistical power of individual experiments to detect typical differences.

Our first objective in this study is to analyze the distribution of effect sizes and statistical power in a large sample of articles using different interventions, showing how they are related to the outcome of statistical significance tests. Next, we will study whether these two measures are correlated, in order to look for evidence of publication bias and effect size inflation. We will also correlate effect sizes and variances with different aspects of experimental design, such as species, sex and type of conditioning, as well as with indicators of risk of bias. To inquire whether effect size and power are taken into consideration by authors when interpreting findings, we will evaluate whether they correlate with effect size inferences made by readers based on textual descriptions of results in the articles. Finally, we will analyze whether mean effect size and power correlate with article-level metrics, such as number of citations and impact factor of the publishing journal, to explore how they influence the publication of results.

## Results

### Article search and inclusion

As previously described in a protocol published in advance of full data collection [20], we performed a PubMed search for fear conditioning articles published online in 2013. The search process (**Fig. 1**) yielded 400 search hits, of which 386 were original articles that were included if they fulfilled pre-established criteria (see Methods). Two investigators examined all included articles, and agreement for exclusions measured on a double-screened sample of 40 articles was 95%. This led to a final sample of 122 articles and 410 experiments, used to build the database provided as **Supplementary Data**.

**Figure 1.**
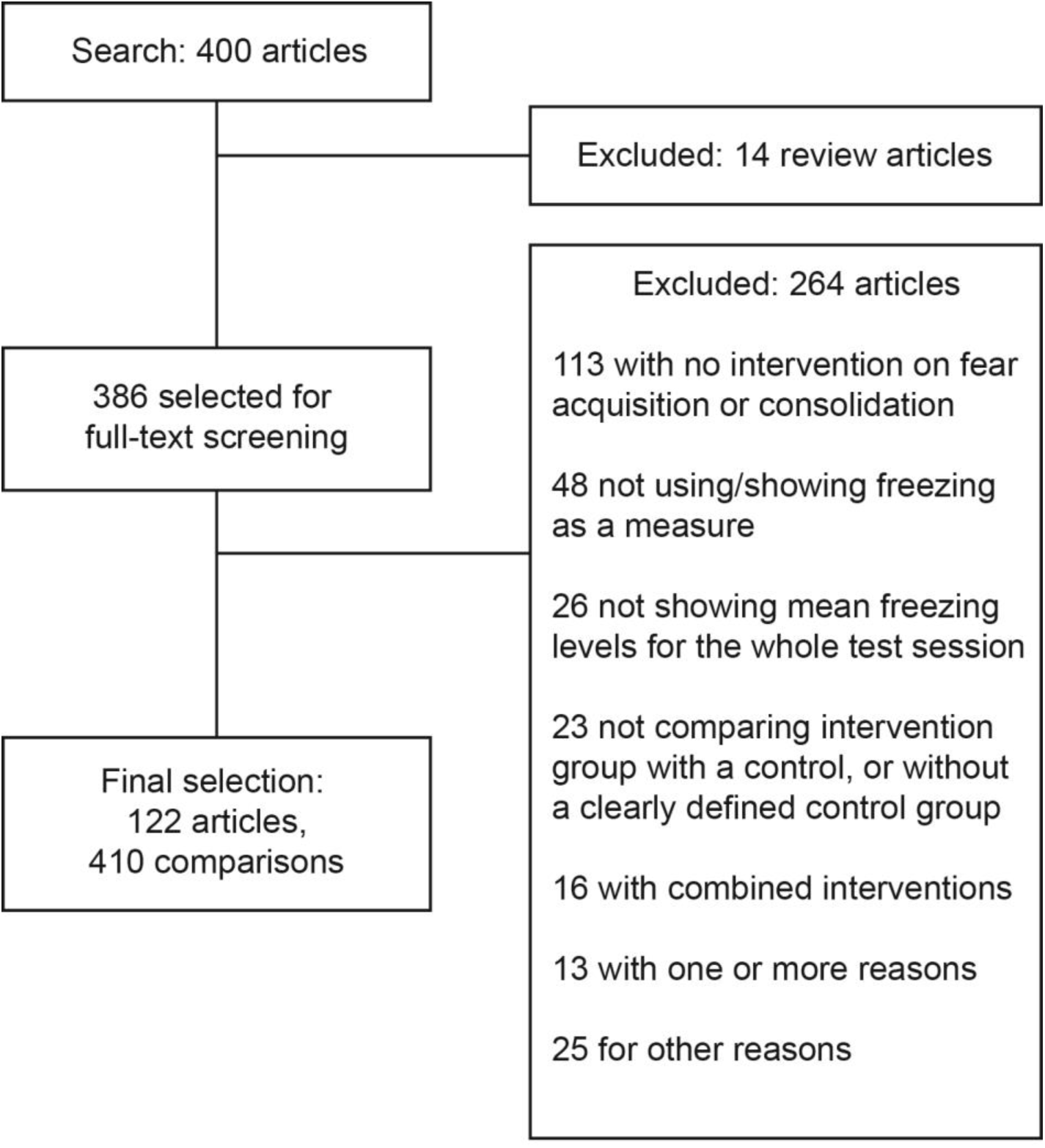
Study flow diagram. Our PubMed search yielded 400 results, of which 14 were excluded based on initial screening of titles and abstracts and 386 were selected for full-text analysis. This led to the inclusion of 122 articles, containing a total of 410 comparisons (i.e. individual experiments). The main reasons for exclusion are listed in the figure, in compliance with the PRISMA statement [21].

### Distribution of effect sizes among experiments

For each experiment, we initially calculated effect size as the relative difference (i.e. percentage of change) in the freezing levels of treated groups when compared to controls. As shown in **Fig. 2A**, this leads interventions that enhance memory acquisition (i.e. those in which freezing is significantly higher in the treated group) to have larger effect sizes than those that impair it (i.e. those in which freezing is significantly lower in the treated group) due to an asymmetry that is inherent to ratios. To account for this and make effect sizes comparable between both types of interventions, we used a normalized effect size, with difference expressed as a percentage of the highest freezing value between groups (**Fig. 2B**) [11].

**Figure 2.**
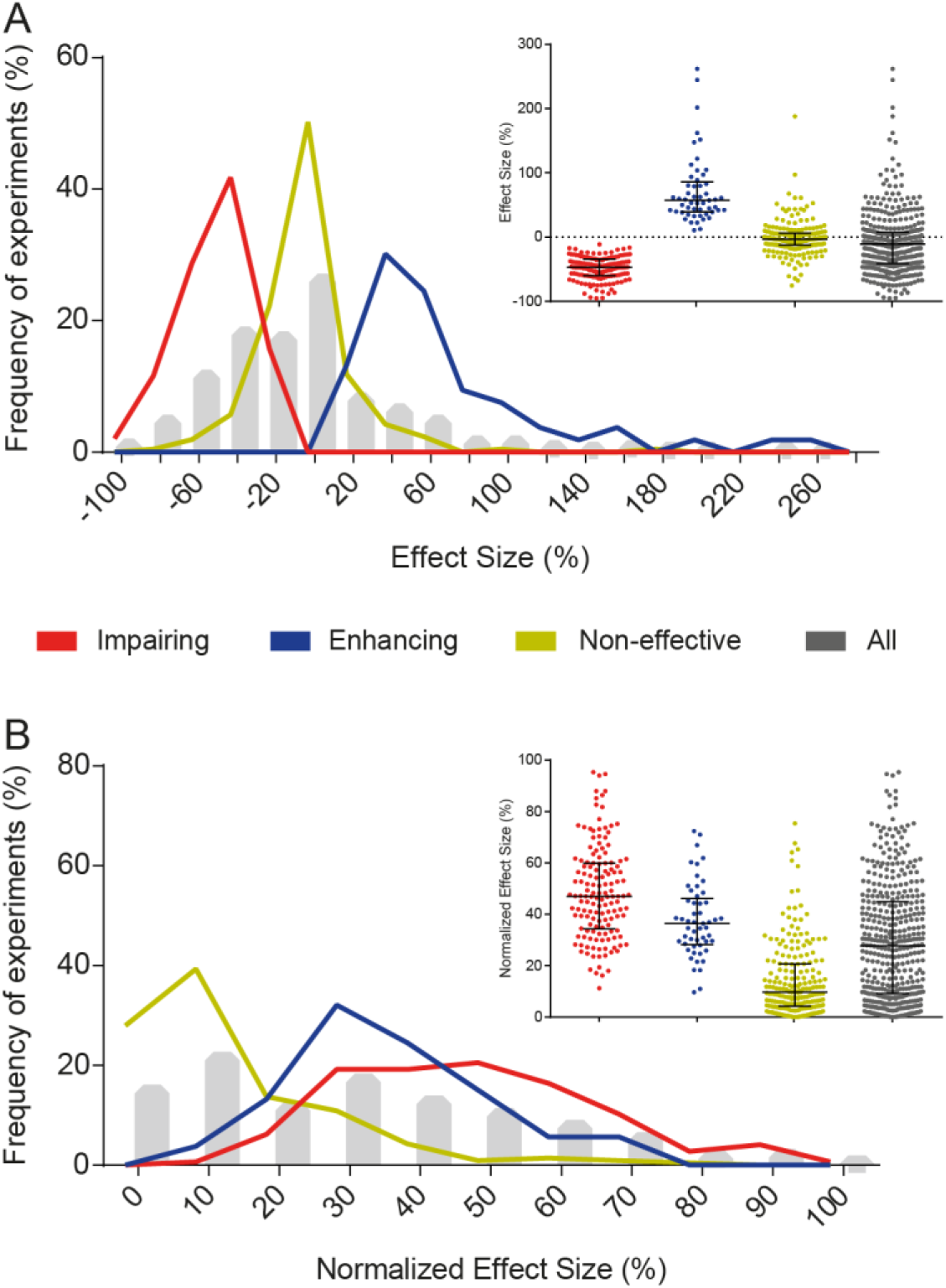
Distribution of effect sizes. (A) Distribution of effect sizes, calculated as % of control group freezing. Interventions were divided into memory-impairing (-48.6 ± 18.1%, n=146), memory-enhancing (71.6 ± 53.2%, n=53) or non-effective (-1.8 ± 26.2%, n=211) for visualization purposes, according to the statistical significance of the comparison performed in the article. Additionally, the whole sample of experiments is shown in grey (-9.0 ± 47.5% [-13.6 to -4.4], n=410). Values are expressed as mean ± SD [95% confidence interval]. Lines and whiskers in the inset express median and interquartile interval. (B) Distribution of normalized effect sizes, calculated as % of the group with the highest mean (i.e. control group for memory-impairing interventions, or treated group for memory-enhancing interventions).

Use of absolute differences in freezing instead of relative ones led to similar, but more constrained distributions (**S1 Fig.**) due to mathematical limits on absolute differences. Freezing levels in the reference group correlated negatively with relative effect size and pooled coefficient of variation (i.e. the ratio between the sample size-weighted pooled SD and the pooled mean); however, normalization by the highest-freezing group reduced this effect (**S2 Fig. A-C**). Absolute effect size, on the contrary, showed a positive correlation with freezing levels in the control or highest-freezing group (**S2 Fig. D-F**). We also calculated effect sizes as standardized mean differences (i.e Cohen’s d, **S3 Fig.**), but chose to use relative percentages throughout the study, as they are more closely related to the way results are expressed in articles.

All 410 experiments combined had a mean normalized effect size of 29.5 ± 22.4% (mean ± SD; 95% CI [27.4 to 31.7]). When results were divided according to the statistical comparison originally performed in the article, mean normalized effect size was 48.6 ± 18.1% for memory-impairing interventions, 37.6 ± 14.2% for memory-enhancing interventions and 14.4 ± 14.2% for non-effective interventions – i.e. those in which a significant difference between groups was not found. This does not imply that data in each of these groups represents effects coming from a different distribution, or that significant and non-significant results correspond to true positive or true negative effects. On the contrary, each group likely represents a mixture of heterogeneous effect size distributions, as sampling error and lack of statistical power can directly impact the chances of a result achieving statistical significance. Distribution of mean effect sizes at the article level showed similar results to those found at the level of experiments (**S4 Fig.**).

The distribution of effect sizes shows that the vast majority of memory-impairing interventions cause partial reductions in learning, leaving the treated group with residual freezing levels that are higher than those of a non-conditioned animal. In fact, in all 35 memory-impairing experiments in which pre-conditioning freezing levels were shown for the treated group, these were lower than those observed in the test session – with p values below 0.05 in 25 (78%) out of the 32 cases in which there was enough information for us to perform an unpaired *t* test between sessions **(S5 Fig.)**. It is also worth noting that 26.5% of non-significant experiments had an effect size greater than 20%, suggesting that these experiments might have been underpowered. With this in mind, we went on to evaluate the distribution of statistical power among studies.

### Distribution of statistical power among experiments

For analyzing statistical power, we first sought to evaluate the distribution of sample sizes and coefficients of variation (both of which are determinants of power). As shown in **Fig. 3A**, most experiments had mean sample sizes between 8 and 12 animals/group, and this distribution did not vary between enhancing, impairing and non-effective interventions. On the other hand, higher coefficients of variation were more frequent among non-effective interventions (**Fig. 3B**). This difference was partly explained by freezing levels in the reference group – which correlated negatively with coefficients of variation (**S2 Fig. G-I**) and were lower on average for non-significant experiments (49.3% vs. 52.9% in memory-impairing and 61.3% in memory-enhancing experiments).

**Figure 3.**
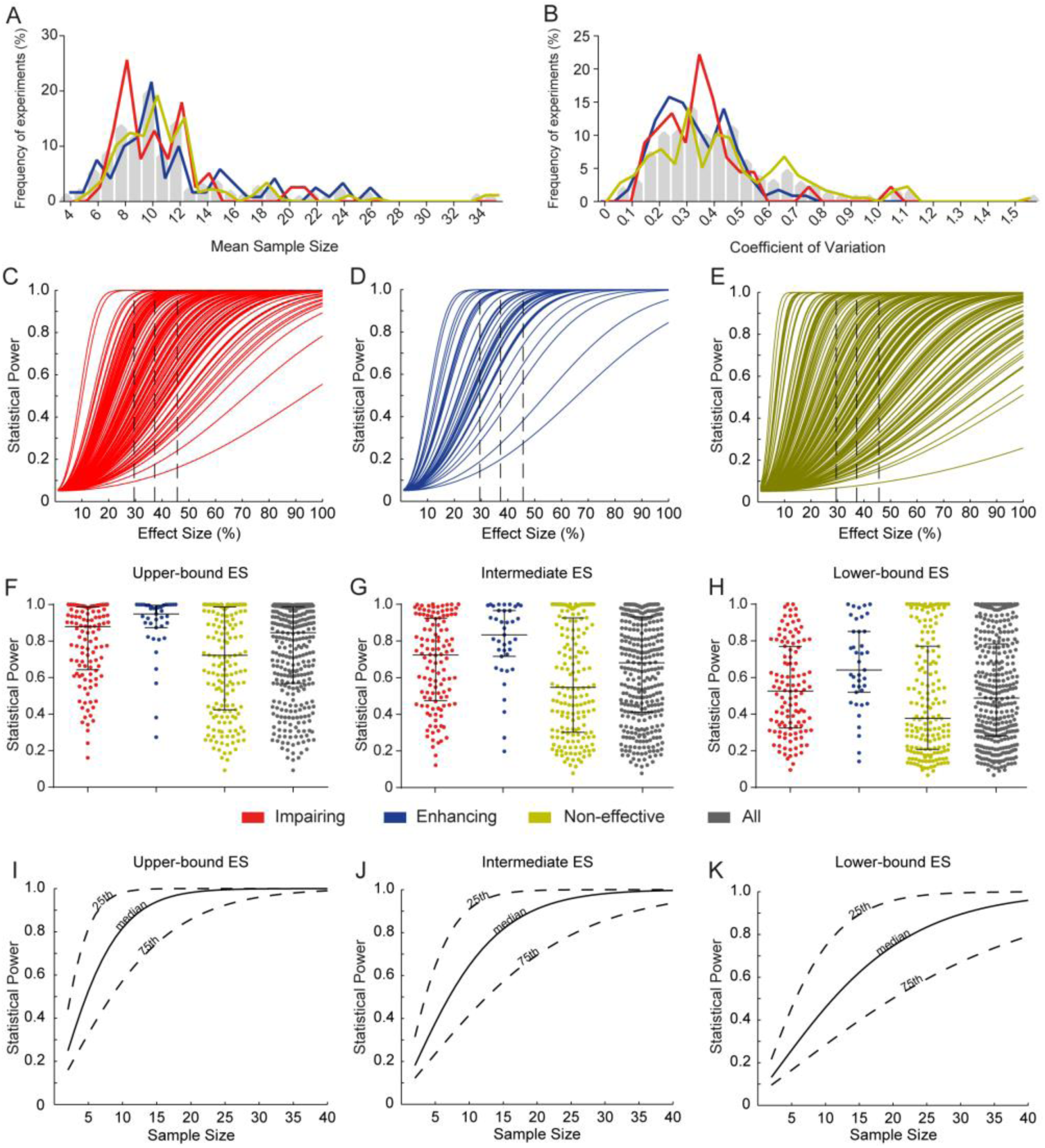
Distribution of sample size, variation and statistical power. (A) Distribution of mean sample size between groups. Gray bars show the distribution of the whole sample, while colored lines show distributions of impairing (n=120), enhancing (n=39) and non-significant (n=177) experiments separately for visualization purposes. (B) Distribution of coefficients of variation (pooled standard deviation/pooled mean) for each type of experiment. (C) Distribution of statistical power for memory-impairing interventions: based on each experiment’s variance and sample size, power varies according to the difference to be detected for α=0.05. Dashed lines show the three effect sizes used for point estimates of power in F, G and H. (D) Distribution of statistical power for memory-enhancing interventions. (E) Distribution of statistical power for non-effective interventions. (F) Distribution of statistical power to detect the upper-bound effect size of 45.6% (right dashed line on C, D and E) for impairing (red), enhancing (blue), non-significant (yellow) and all (grey) experiments. Lines and whiskers express median and interquartile interval. (G) Distribution of statistical power to detect the intermediate effect size of 37.2% (middle dashed line on C, D and E). (H) Distribution of statistical power to detect the lower-bound effect size of 29.5% (left dashed line on C, D and E). (I) Sample size vs. statistical power to detect the upper-bound effect size of 45.6%. Continuous lines use the 50^th^ percentile of coefficients of variation for calculations, while dotted lines use the 25^th^ and 75^th^ percentiles. (J) Sample size vs. statistical power to detect the intermediate effect size of 37.2%. (K) Sample size vs. statistical power to detect the lower-bound effect size of 29.5%.

Based on each experiment’s variance and sample size, we built power curves to show how power varies according to the difference to be detected at α=0.05 for each individual experiment (**Fig. 3C-E**). To detect the mean effect size of 45.6% found for nominally effective interventions (i.e. those leading to statistically significant differences between groups), mean statistical power in our sample was 0.75 ± 0.26; 95% CI [0.72 - 0.78] (**Fig. 3F**). This estimate, however, is an optimistic, upper-bound calculation of the typical effect size of biologically effective interventions (from here on referred to as “upper-bound ES”): as only large effects will be detected by underpowered studies, basing calculations only on significant results leads to effect size inflation [14]. A more realistic estimate of effect size was obtained based only on experiments that achieved statistical power above 0.95 (n=60) in the first analysis (and are thus less subject to effect size inflation), leading to a mean effect size of 37.2%. Predictably, mean statistical power to detect this difference (“intermediate ES”, **Fig. 3G**) fell to 0.65 ± 0.28 [0.62 - 0.68]. Using the mean effect size of all experiments (“lower-bound ES”, 29.5%) led to an even lower power of 0.52 ± 0.29 [0.49 - 0.56] (**Fig. 3H**), although this estimate of a typical effect size is likely pessimistic, as it probably includes many true negative effects.

Interestingly, using mean absolute differences instead of relative ones to calculate statistical power led to a smaller number of experiments with very low power (**S6 Fig.**). This suggests that some of the underpowered experiments in the first analysis had low freezing levels in the reference group, as in this case even large relative differences will still be small when expressed in absolute terms for statistical analysis. Also of note is that, if one uses Cohen’s traditional definitions of small (d=0.2), medium (d=0.5) and large (d=0.8) effect sizes [22] as the basis for calculations, mean power is 0.07 ± 0.01, 0.21 ± 0.07 and 0.44 ± 0.13, respectively (**S7 Fig.**). These much lower estimates reflect the fact that effect sizes are typically much larger in rodent fear conditioning than in psychology experiments, for which this arbitrary classification was originally devised, and suggests that it might not be applicable to other fields of science.

A practical application of these power curves is that we were able to calculate the necessary sample size to achieve desired power for each effect size estimate, considering the median coefficient of variation (as well as the 25^th^ and 75^th^ quartiles) of experiments in our sample (**Fig. 3I-K**). Thus, for an experiment with typical variation, around 15 animals per group are needed to achieve 80% power to detect our ‘intermediate effect size’ of 37.2%, which we consider our more realistic estimate for a typical effect size in the field. Nevertheless, only 12.2% of comparisons in our sample had a sample size of 15 or above in each experimental group, suggesting that such calculations are seldom performed.

We also analyzed the distributions of statistical power at the level of articles instead of individual experiments. Results for these analyses are shown in **S8 Fig.**, and are generally similar to those obtained for the experiment-level analysis, except that the long tail of non-significant experiments with large coefficients of variation is not observed. This suggests that experiments with large variation and low power are frequently found alongside others with adequate power within the same articles. It is unclear, however, whether this means that the low power of some experiments is a consequence of random fluctuations of experimental variance, or if these experiments use protocols that lead to larger coefficients of variation – for example, by generating lower mean levels of freezing (see **S2 Fig.**).

### Correlation between effect sizes and statistical power/sample size

We next sought to correlate normalized effect size with sample size and statistical power for each experiment. The presence of a negative correlation between these variables has been considered an indirect measure of publication bias [23], as articles with low power or sample size will be subject to effect size inflation caused by selective reporting of significant findings [24]. In our analysis, no correlation was found between effect size and sample size (**Fig. 4A**, r=0.0007, p=0.99); on the other hand, a positive correlation between effect size and coefficient of variation was observed (**Fig. 4B**, r=0.37, p<0.0001). Part of this correlation was mediated by the association of both variables with freezing levels (**S2 Fig.**), but the correlation remained significant after adjustment for this variable (r=0.32, p<0.001).

**Figure 4.**
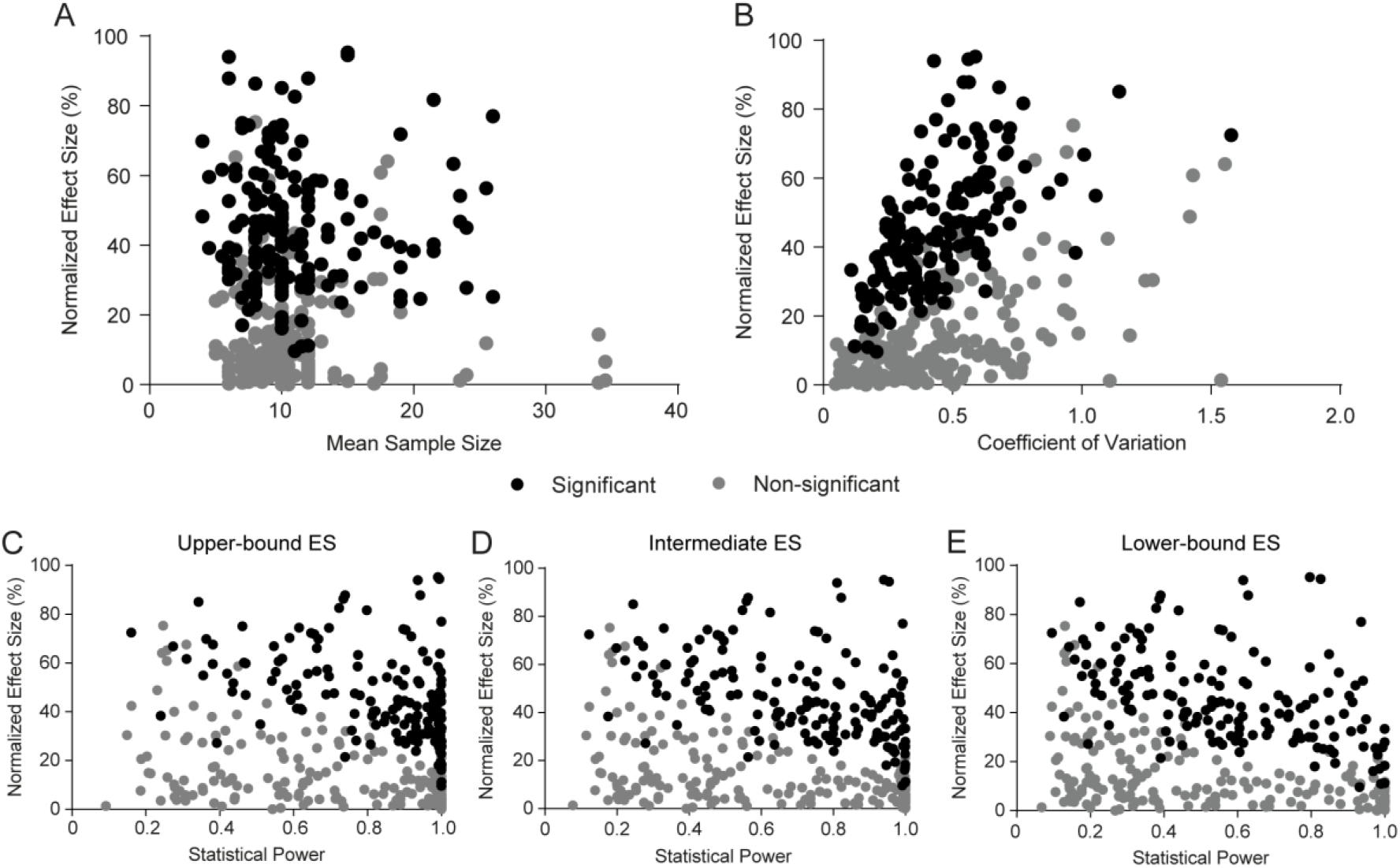
Correlations between effect size, variation and statistical power. (A) Correlation between normalized effect size and mean sample size. No correlation is found (r=0.0007, p=0.99; r=-0.26, p=0.64 after adjustment), although sample size variation is limited. (B) Correlation between normalized effect size and coefficient of variation. Correlation of the whole sample of experiments yields r=0.37, p<0.0001* (n=336; r=0.32, p<0.001 after adjustment for freezing levels). (C) Correlation between normalized effect size and statistical power based on upper-bound effect size of 45.6%. Correlation of the whole sample of experiments yields r=-0.12, p=0.03 (r=0.11, p=0.84 after adjustment for freezing levels), but distribution is skewed due to a ceiling effect on power. (D) Correlation between normalized effect size and statistical power based on intermediate effect size of 37.2%; r=-0.16, p=0.003* (r=-0.16, p=0.48 after adjustment). Correlation between normalized effect size and statistical power based on lower-bound effect size of 29.5%; r=-0.21, p<0.0001* (r=-0.1, p=0.06 after adjustment). Asterisks indicate significant results according to Holm-Sidak correction for 28 experiment-level correlations.

Because of this, negative correlations between effect size and power were observed for the three effect size estimates used (**Figs. 4C-E**), although they were larger for the lower-bound estimate (**Fig. 4E**, r=-0.21, p<0.0001) than for the intermediate (**Fig. 4D**, r=-0.16, p=0.003) and upper-bound (**Fig. 4C**, r=-0.12, p=0.03) ones due to a ceiling effect on power. This negative correlation is observed even when power is calculated based on absolute differences (**S9 Fig.**), for which the correlation between coefficients of variation and reference freezing levels is in the opposite direction of that observed with relative differences (see **S2 Fig.**). This strongly suggests that the correlation represents a real phenomenon related to publication bias and/or effect size inflation, and is not merely due to the correlation of both variables with freezing levels. A correlation between effect size and power is also observed when both are calculated on the basis of standardized mean differences (i.e. Cohen’s d) **(S10 Fig.**). In this case, the line separating significant and non-significant results for a given sample size is clearer, as significance is more directly related to standardized mean differences. Expressing effect sizes in Cohen’s d also makes effect size inflation in experiments with low sample size and power more obvious.

Interestingly, the correlation between effect size and power was driven by a scarcity of experiments with large effect size and high power. This raises the possibility that truly large effects are unusual in fear conditioning, and that some of the large effect sizes among low-powered experiments in our sample are inflated. On the other hand, a pattern classically suggesting publication bias – i.e. a scarcity of low-powered experiments with small effects [23] – is not observed. It should be noted, however, that our analysis focused on individual experiments within articles, meaning that non-significant results were usually presented alongside other experiments with significant differences; thus, this analysis does not allow us to assess publication bias at the level of articles.

### Effects of methodological variables on the distributions of effect sizes and coefficients of variation

We next examined whether the distributions of effect sizes and coefficients of variation were influenced by type of conditioning, species or sex of the animals (**Fig. 5**). Mean normalized effect size was slightly larger in contextual than in cued fear conditioning (33.2% vs. 24.4%, Student’s t test p<0.0001) and markedly larger in males than in females (30.3% vs. 18.9% vs. 34.2% for experiments using both, one-way ANOVA, p=0.004), but roughly equivalent between mice and rats (29.8% vs. 29.1%, p=0.76). Coefficients of variation were higher in contextual conditioning (0.51 vs. 0.41, Student’s t test p=0.001), in experiments using animals of both sexes (0.62 vs. 0.44 in males and 0.41 in females, one-way ANOVA, p <0.0001), and in those using mice (0.50 vs. 0.42, Student’s t test, p=0.008), although the latter difference was not statistically significant after correction for multiple comparisons. All of these associations should be considered correlational and not causal, as specific types of conditioning or animals of a particular species or sex might be more frequently used for testing interventions with particularly high or low effect sizes. Also of note is the fact that experiments on males were 7.7 times more common than those on females in our sample (277 vs. 36), indicating a strong preference of researchers for using male animals.

**Figure 5.**
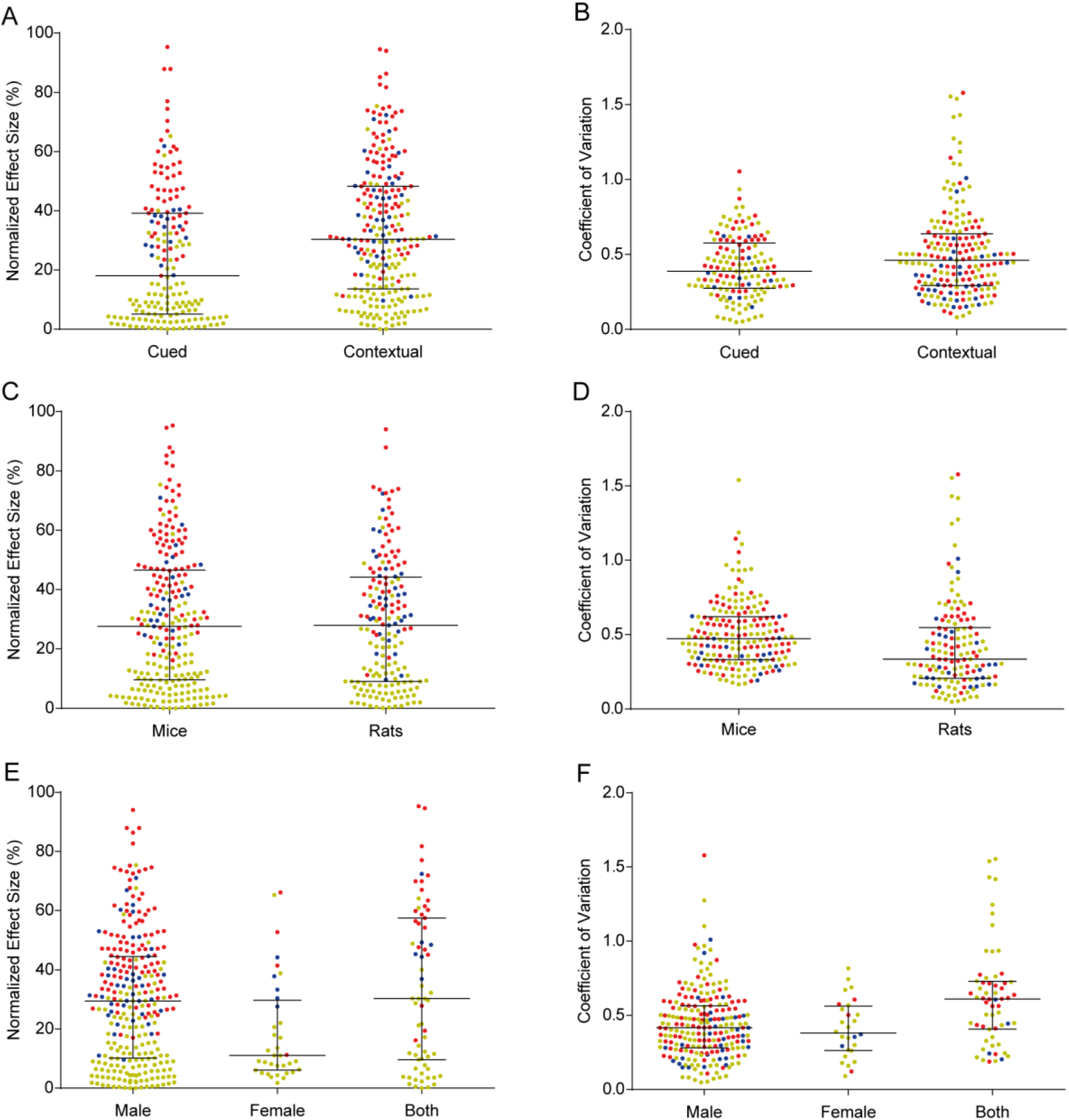
Effect sizes and coefficients of variation across different protocols, species and sexes. Colors indicate memory-enhancing (red), memory-impairing (blue) or non-effective (yellow) experiments, all of which are pooled for analysis. Lines and whiskers express median and interquartile interval. (A) Distribution of effect sizes across cued (n=171) and contextual (n=239) conditioning protocols. Student’s t test, p<0.0001*. (B) Coefficients of variation across cued (n=145) and contextual (n=191) conditioning protocols. Student’s t test, p=0.001*. (C) Distribution of effect sizes across experiments using mice (n=237) or rats (n=173). Student’s t test, p=0.76. (D) Coefficients of variation across experiments using mice (n=193) or rats (n=143). Student’s t test, p=0.008. (E) Distribution of effect sizes across experiments using male (n=277), female (n=36) or both (n=67) sexes. One-way ANOVA, p=0.004*; Tukey’s post-hoc test, male vs. female p=0.01, male vs. both p=0.40, female vs. both p=0.003. 30 experiments were excluded from this analysis for not stating the sex of animals. (F) Coefficients of variation across experiments using male (n=233), female (n=28) or both (n=60) sexes. One-way ANOVA, p<0.0001*; Tukey’s test, male vs. female p=0.85, male vs. both p<0.0001, female vs. both p=0.0006. For coefficient of variation analyses, 74 experiments were excluded due to lack of information on sample size for individual groups. Asterisks indicate significant results according to Holm-Sidak correction for 14 experiment-level comparisons.

We also examined whether effect sizes and coefficients of variation differed systematically according to the type, timing or anatomical site of intervention (**S11 Fig.**). Effect sizes did not differ significantly between surgical, pharmacological, genetic and behavioral interventions (38.7% vs. 28.1% vs. 30.5% vs. 25.8% one-way ANOVA, p=0.12), although there was a trend for greater effects with surgical interventions (which were uncommon in our sample). No differences were found between the mean effect sizes of systemic and intracerebral interventions (28.7% vs. 30.3%, Student’s t test, p=0.45) or between those of pre- and post-training interventions (30.5% vs. 25.4%, Student’s t test, p=0.07), although pre-training interventions had slightly higher coefficients of variation (0.49 vs 0.37, Student’s t test p=0.0015). Coefficients of variation did not differ significantly between surgical, pharmacological, genetic and behavioral interventions (0.41 vs. 0.43 vs. 0.50 vs. 0.50, one-way ANOVA p=0.08) or between systemic and intracerebral interventions (0.49 vs. 0.45, Student’s t test p=0.15). Once again, these differences can only be considered correlational and not causal.

### Risk of bias indicators and their relationship with effect size and power

As previous studies have shown that measures to reduce risk of bias are not widely reported in animal research [12,13], we investigated the prevalence of these measures in our sample of fear conditioning articles, and evaluated whether they were correlated with effect sizes or power. **Table 1** shows the percentage of articles reporting 7 items thought to reduce risk of bias in animal studies, adapted and expanded from the CAMARADES checklist [25]. Although some items were reported in most articles (statement of compliance with animal regulations, adequate description of sample size, blinding), others were virtually inexistent, such as the presence of a sample size calculation (1 article) and compliance with the ARRIVE guidelines [26] (0 articles). Contrary to previous reports in other areas [27–30], however, no significant association was found between reporting of these indicators and either the percentage of significant experiments, the mean effect size of effective interventions or the mean statistical power of experiments in our sample (**S12 Fig.**). The region of origin of the article also had no correlation with either of these variables (**S13 Fig.**). Nevertheless, it should be noted that this analysis used only experiments on fear conditioning acquisition or consolidation, which were not necessarily the only results or the main findings presented in these articles. Thus, it is possible that other results in the article might have shown higher correlation with risk of bias indicators.

**Table 1.**
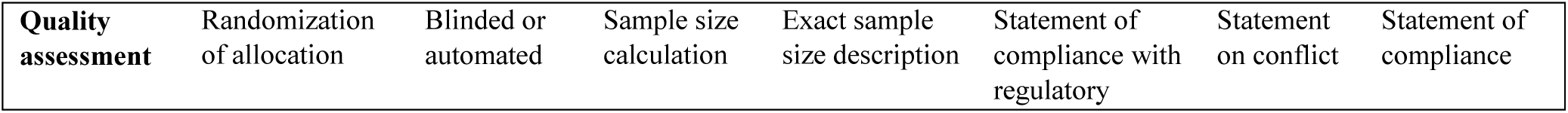
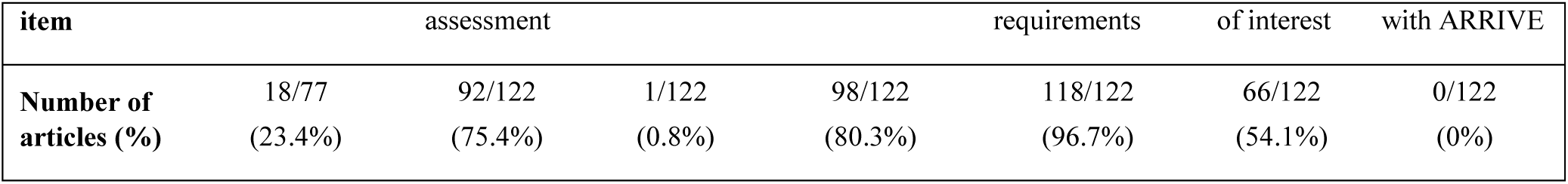
Number of articles including quality assessment items. Percentages were calculated using all 122 articles, except in the case of randomization, which was calculated based on 77 articles, as it is not applicable to genetic interventions. In the case of blinding, 68 articles used automated analysis and 24 used blinded observers, totaling 92 articles scored for this item.

### Correlation between effect sizes/statistical power and description of results

Given the wide distribution of effect sizes and statistical power in the literature on fear conditioning learning, we tried to determine whether these were taken into account by authors when describing results in the text. For each included comparison, we extracted the words or phrases describing the results of that experiment in the text or figure legends, and asked 14 behavioral neuroscience researchers to classify them according to the implicit information they contained about effect size. For comparisons with significant differences, terms were to be classified as implying strong (i.e. large effect size) or weak (i.e. small effect size) effects, or as neutral terms (i.e. those from which effect size could not be deduced). For non-significant differences, terms were to be classified as implying similarity between groups, as suggesting a trend towards difference, or as neutral terms (i.e. those from which the presence or absence of a trend could not be deduced). From the average of these classifications, we defined a score for each term (**S1 and S2 Tables**) and correlated these scores with the actual effect size and statistical power of experiments.

Agreement between researchers over classification was low, especially for terms describing significant differences: single measures intraclass correlation coefficients (reflecting the reliability of individual researchers when compared to the whole sample) were 0.234 for significant interventions and 0.597 for non-significant ones, while average measures coefficients (reflecting the aggregated reliability of the sample) were 0.839 and 0.962, respectively. This, along with a trend for the use of terms with little effect size information (“increase”, “decrease”, “significantly more”, “significantly less”, etc.), led most terms describing effective interventions to receive intermediate scores approaching 1 (i.e. neutral). For these interventions, no correlations were observed between this score and either effect size (r=-0.05, p=0.48) or statistical power (r=0.03, p=0.73) (**Fig. 6A and 6B**). For non-effective interventions, a significant correlation between description score and effect size was observed (**Fig 6C**, r=0.28, p=0.0002), as larger effect sizes were associated with terms indicating a trend for difference. Still, no correlation was observed between textual descriptions of results and power (**Fig 6D**, r=0.03, p=0.74). Moreover, statistical power was rarely mentioned in the textual description of results – the term “power” was used in this context in only 4 articles– suggesting that it is largely ignored when discussing findings, as shown in other areas of research [31].

**Figure 6.**
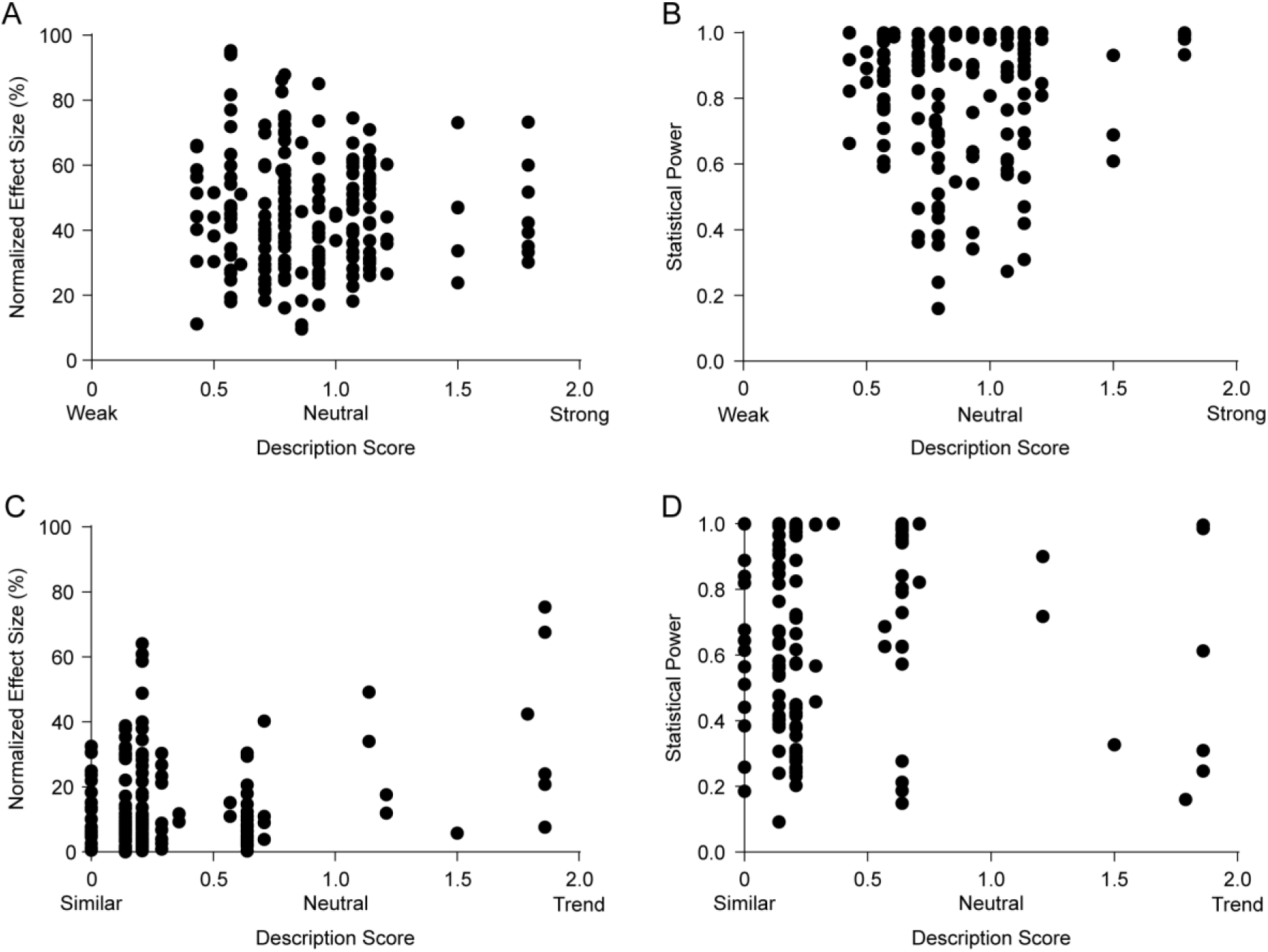
Correlation between description of results and effect size/statistical power. Description scores refer to the mean score given by 14 neuroscience researchers who rated terms as “weak” (0), “neutral” (1) or “strong” (2) in the case of those describing significant differences, or as “similar” (0), “neutral” (1) or “trend” (2) in the case of those describing non-significant ones. (A) Correlation between normalized effect size and description score for significant results. r=-0.05, p=0.48 (n=195). (B) Correlation between statistical power and description score for significant results. r=0.03, p=0.73 (n=155). (C) Correlation between normalized effect size and description score for non-significant results. r=0.28, p=0.0002* (n=174). (D) Correlation between upper-bound estimate of statistical power and description score for non-significant results. r=0.03, p=0.74 (n=146). Asterisk indicates significant result according to Holm-Sidak correction for 28 experiment-level correlations.

### Correlations of effect size, power and study quality with article citations

Finally, we investigated whether the percentage of significant experiments reported in each article, mean effect size for effective interventions, mean statistical power or a composite study quality score (aggregating the 7 risk of bias indicators described in **Table 1**) correlated with article impact, as measured by the number of citations (**Fig. 7**) and the impact factor of the publishing journal (**S14 Fig.**). None of the correlations was significant after adjustment for multiple comparisons, although a weak positive correlation was observed between study quality score and impact factor (r=0.22, p=0.01), driven by associations of higher impact factors with blinding (Student’s t test with Welch’s correction, p=0.0001), conflict of interest reporting (Student’s t test with Welch’s correction, p=0.03) and exact sample size description (Student’s t test, p=0.03). It should be noted that the distribution of impact factors and citations is heavily skewed, limiting the use of linear correlations as planned in the original protocol – nevertheless, exploratory non-parametric analysis of the data confirmed the lack of significance of correlations. Once again, our data refers only to experiments on fear conditioning acquisition or consolidation – therefore, other data in the articles could feasibly account for the variation in impact factor and citations.

**Figure 7.**
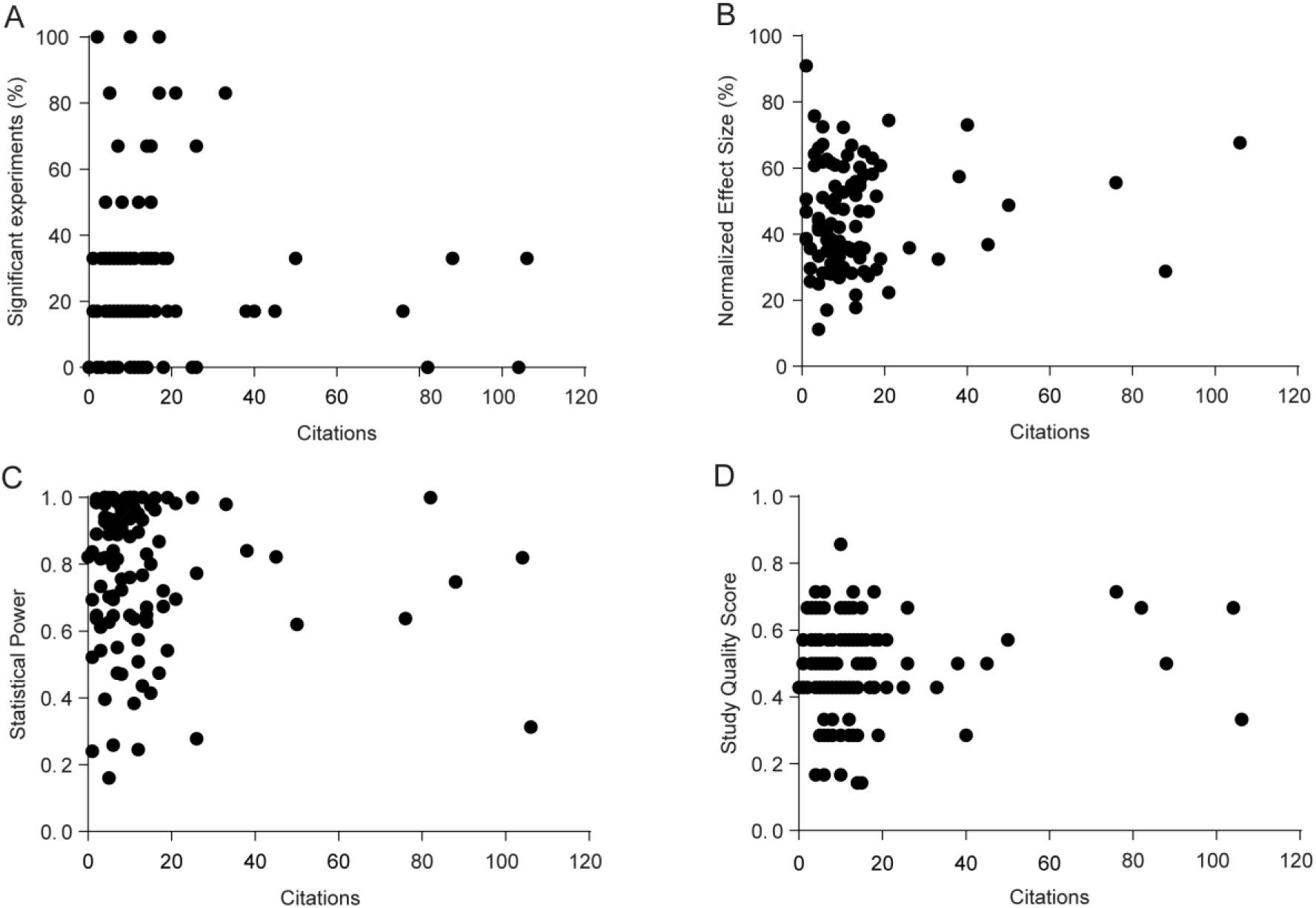
Correlation between citations and percentage of significant experiments, effect size and statistical power. Citations were obtained for all articles on August 26^th,^ 2016. (A) Correlation between % of significant results per article and citations. r=-0.03, p=0.75 (n=121). (B) Correlation between mean normalized effect size of effective interventions and citations. r=0.097, p=0.34 (n=98). (C) Correlation between mean statistical power (upper-bound estimate) and citations. r=-0.08, p=0.40 (n=104). (D) Correlation between study quality score and citations. r=0.09, p=0.31 (n=121). According to Holm-Sidak correction for 8 article-level correlations, none is significant.

## Discussion

In light of the low reproducibility of published studies in various fields of biomedical research [32–34] which is thought by many to be a consequence of low statistical power and excessive reliance on significance tests [8,16] calls have been made to report effect sizes and confidence intervals alongside or in place of p values [4,6,7,9] and to increase statistical power [14,31,35]. However, it is unclear whether these proposals have had much impact on most fields of basic science. We have taken one particular memory task in rodents, in which outcomes and effect sizes are described in a standardized way and are thus comparable across studies, in order to analyze how these two concepts are dealt with in the study of fear learning.

Our first main finding is that most amnestic interventions targeting fear acquisition or consolidation cause partial effects, with residual freezing remaining significantly above pre-conditioning levels in 78% of the experiments with available data. Moreover, most of the large effect sizes in our sample were found in underpowered studies, suggesting that they could represent inflated estimates [24]. This is not necessarily unexpected: as fear memories depend on a well distributed network, both anatomically and molecularly [19], it seems natural that most interventions directed at a specific site or pharmacological target will modulate learning rather than fully block it. This creates a problem, however, when effect sizes are not considered in the analysis of experiments, as it is not possible to differentiate essential mechanisms of memory formation from modulatory influences on the basis of statistical significance alone. This can lead to a situation in which accumulating evidence, even if correct, can confuse rather than advance understanding, as has been suggested to occur in fields such as long-term potentiation [15] and apoptosis [36].

Matters are complicated further by the possibility that many of these findings are false positives and/or false negatives. The prevalence of both in relation to true positives and negatives depends crucially on statistical power, which in turn depends on sample size. Calculating the actual power of published experiments is challenging, as the difference used for the calculations should not be based on the observed results – which leads power analysis to become circular [18,37]. Thus, statistical power depend on expected effect sizes, which are arbitrary by nature – although they can sometimes be estimated from meta-analyses [14], which were not performed in this study due to the large variety of heterogeneous interventions. However, by considering the mean effect size for well-powered experiments in our sample, we arrived at an estimate of around 37.2% that might be considered “typical” for a published experiment with an intervention affecting fear conditioning acquisition or consolidation. Using the sample size and variation for each experiment, we found mean statistical power to detect this effect size to be 65% in our sample.

As sample size calculations are exceedingly rare, and insufficient power seems to be the norm in other fields of neuroscience as well [14,18], it is quite possible that classically used sample sizes in behavioral neuroscience (and perhaps in other fields of basic science) might thus be insufficient. Considering median variances and our intermediate effect size estimate, the ideal sample size to achieve 80% power would be around 15 animals per group. This number, however, was reached in only 12.2% of cases in our sample, as most experiments had sample sizes of 8 to 12, informally considered to be standard in the field. This seems to confirm recent models suggesting that current incentives in science favor the publication of underpowered studies [16,38], although they could also be due to restrictions on animal use imposed by ethical regulations. That said, average power in our sample for typical effect sizes was higher than those described in other areas of neuroscience by Button et al. [14]; however, this could reflect the fact that effect sizes in their study were calculated by meta-analysis, and might be smaller than those derived by our method of estimation, or underestimated due to the inclusion of negative results [18]. One should also note that the abovementioned power estimates were found to vary widely across subfields of neuroscience [18] – in this sense, the power distribution of fear conditioning studies seems to resemble those found for psychology and neurochemistry studies, in which a reasonable number of well-powered studies coexist with underpowered ones.

On the other hand, our statistical power to detect Cohen’s definitions of small, medium and large effects [22] were even lower than those recently reported in cognitive neuroscience studies by Szucs and Ioannidis (2017). That said, our data provides a strong cautionary note against the use of these arbitrary definitions, originally devised for psychology studies, in calculations of statistical power, as 88.7% of statistically significant experiments (or 48.2% of the whole sample) fell into the “large” category of Cohen’s original proposal. This suggests that laboratory studies in rodents have larger effects than those found in human psychology (an unsurprising finding, given the greater invasiveness of the interventions), as has also been found in meta-analyses studying similar treatments in laboratory animals and humans [39], demonstrating that what constitutes a small or large effect can vary between different fields of science.

An old-established truism in the behavioral neuroscience field – as well as in other fields of basic science – is that experiments in females tend to yield more variable results due to estrous cycle variations [40]. However, at least in our analysis, coefficients of variation were similar between experiments in males and females (and predictably higher in experiments using both), as has been found in other areas of science [41,42] suggesting that this belief is false. Nevertheless, adherence to it likely accounts for the vast preponderance of experiments on male animals, which were nearly 8 times more common than those in females in our sample – a sex bias greater than those described for most fields [43] although smaller than that recently reported for rodent models of anxiety [44]. Previous work in clinical [45] and preclinical [40,43] data has pointed out the drawbacks of concentrating experiments in male populations. However, despite calls for increasing the number of studies on females [46] this problem remains strikingly present in the fear learning field.

Concerning risk of bias indicators, the prevalence found in our sample was roughly similar to previous reports on animal studies for randomization and conflict of interest reporting [13] but were distinctly higher for blinded assessment of outcome, largely because 59% of articles used automated software to measure freezing, which we considered to be equivalent to blinded assessment. If one considers only articles reporting manual scoring of freezing, however, blinding was reported in 57% of cases, which is still higher than most areas of preclinical science [13]. As described previously in many fields [12,13,31] sample size calculations were almost non-existent, which helps to explain why many experiments are underpowered. Interestingly, although we analyzed a sample of papers published 3 years after the ARRIVE guidelines they were not mentioned in any of the articles, suggesting that their impact, at least in the field of behavioral neuroscience, was still rather limited at this time.

Contrary to previous studies, however, we did not detect an impact of these risk of bias indicators on article-level measures such as percentage of fear conditioning learning experiments with significant results, mean effect size of significant experiments and mean statistical power. This could mean that, compared to preclinical studies, bias towards positive results is lower in studies on fear learning. However, it seems more likely that, as we selected particular experiments within papers containing other results, we were not as likely to detect effects of bias on article-level measures. As basic science articles typically contain numerous results, it is perhaps less likely that all comparisons will be subject to bias towards positive findings. Moreover, the experiments in our sample probably included negative controls for other findings, which might have been expected to yield non-significant results. Thus, although our results do not indicate an impact of bias on article-level results, they should not be taken as evidence that this does not occur.

The same reasoning applies for the evaluation of publication bias, in which the experiments we analyzed could have been published along with positive ones. Nevertheless, we were still able to detect a negative correlation between effect size and statistical power, suggesting effect size inflation due to low statistical power to be present in studies on fear conditioning learning. Although the pattern we detected was less suggestive of actual publication bias, our capability to detect it was likely smaller due to the choice to use experiments within articles. Other methods to detect publication bias, such as the Ioannidis excess significance test [47] and the use of p-value distributions [48–50] were also considered, but found to be inappropriate for use with our methodology (in the first case due to the absence of a meta-analytic effect estimate, and in the second because exact p values were infrequently provided in articles).

One of the most interesting findings of our article was the lack of correlation of effect sizes inferred from textual description of results with the actual effect sizes of significant experiments, as well as with statistical power. Although this suggests that these measures are not usually considered in the interpretation of results, there are caveats to this data. First of all, agreement between what words describe a “strong” or “weak” effect between researchers evaluating them was strikingly low, suggesting that written language is a poor descriptor for quantitative data. Moreover, the fact that most terms used to describe differences were neutral to effect sizes (e.g. “significantly higher”, “significantly lower”, etc.) limited our ability to detect a correlation. That said, the high prevalence of neutral terms by itself is evidence that effect sizes are not usually taken into account when reporting results, as differences tend to be described in the text by their statistical significance only.

This point is especially important to consider in the light of recent calls for basic science to use data synthesis tools such as meta-analysis [11] and formal or informal Bayesian inference [2,8,10,51]. In both of these cases, the incremental effect of each new experiment on researchers’ beliefs on the veracity of a finding is dependent both on the effect size of the result and on its statistical significance. However, even exact p values were uncommonly reported in our sample, with the majority of articles describing p as being above or below a threshold value. This seems to suggest that researchers in the field indeed tend to consider statistical significance as a binary outcome, and might not be quite ready or willing to move towards Bayesian logic, which would require a major paradigm shift in the way results are reported and discussed.

An interesting question is that, if researches in the field indeed were to move away from null-significance hypothesis testing, the concept of statistical power as it is defined today would largely lose its meaning (as it is intrinsically linked to the idea of a significance threshold). Nevertheless, the necessity of adequate sample size for statistical robustness would remain – in this case, not in order to detect significant differences and prevent false-negatives and false-positives, but to estimate effect sizes with adequate precision. The current notion of statistical power to detect a given difference could thus be replaced with a desired confidence interval for the obtained result when performing sample size calculations – a formulation that might be useful in terms of differentiating biologically significant results from irrelevant ones.

Concerning article impact metrics, our results are in line with previous work showing that journal impact factor does not correlate with statistical power [14] or with most risk of bias indicators [13]. Furthermore, we showed that, in articles on fear conditioning, this lack of correlation also occurs for the percentage of significant experiments and the mean effect size for significant differences, and that it extends to citations measured over 2 subsequent years. That said, our article-level analysis was limited by the fact that, for many articles, the included experiments represented a minority of the findings. Moreover, most articles tend to cluster around intermediate impact factors (i.e. between 3 and 6) and relatively low (< 20) citation numbers. Thus, our methodology might not have been adequate to detect correlations between these metrics with article-wide effect size and power estimates.

The choice to focus on a particular type of experiment – in this case, interventions directed at rodent fear conditioning acquisition or consolidation – is both one of the main strengths and the major limitation of our findings. On one hand, it allows us to look at effect sizes that are truly on the same scale, as fear conditioning protocols tend to be reasonably similar across laboratories, and all included experiments described their results using the same metric. Thus, the studied effect sizes are not abstract and have real-life meaning. On the other hand, this decision limits our conclusions to this specific field of science, and also weakens our article-level conclusions, as most articles had only a fraction of their experiments analyzed.

Dealing with multiple experiments using different outcomes presents a major challenge for meta-research in basic science, and all alternatives present limitations. A radically opposite approach of converting all effect sizes in a field to a single metric (e.g. Pearson’s r, Cohen’s d, etc.) has been used by other researchers investigating similar topics in neuroscience and psychology [17,23,31,35]. Although normalizing effect sizes allows one to obtain results from a wider field, it also leads them to be abstract and not as readily understandable by experimental researchers. Moreover, this approach can lead to the aggregation of results from disparate types of experiments for which effect sizes are not in the same scale, leading to important distortions in calculating power for individual experiments. Finally, recent evidence indicates that, even within neuroscience, features such as statistical power have very different distributions across subfields [18], suggesting that surveys of individual areas are likely to be more reliable for studying them.

In our case, studying the concrete scenario of a specific methodology leads to more readily applicable suggestions for experimental researchers, such as the rule-of-thumb recommendation that the average number of animals per group in a fear conditioning experiments to achieve 80% power would be around 15 for typical effect sizes and variances. Our approach also allowed us to detect correlations between results and specific methodological factors (e.g. context vs. cued conditioning, female vs. male animals) that would not be apparent if multiple types of experiments were pooled together. Still, to provide more solid conclusions on the causal influence of these factors on experimental results, even our methodology has too wide a focus, as analyzing multiple interventions limits our possibilities to perform meta-analysis and meta-regression to control for confounding variables. Follow-up studies with more specific aims (i.e. meta-analyses of specific interventions in fear conditioning) are thus warranted to understand the variation between results in the field.

Finally, it is important to note that, while our study has led to some illuminating conclusions, they are inherently limited to the methodology under study. Thus, extrapolating our findings to other types of behavioral studies, not to mention other fields of science, requires data to be collected for each specific subfield. While this might appear herculean at first glance, it is easily achievable if scientists working within specific domains start to design and perform their own systematic reviews. Only through this dissemination of meta-research across different areas of science will we be able to develop solutions that, by respecting the particularities of individual subfields, will be accepted enough to have an impact on research reproducibility.

## Materials and Methods

The full protocol of data selection, extraction and analysis was initially planned on the basis of a pilot analysis of 30 papers, and was registered, reviewed and published ahead of full data extraction [20]. In brief, we searched PubMed for the term “fear conditioning” AND (“learning” OR “consolidation” OR “acquisition”) AND (“mouse” OR “mice” OR “rat” OR “rats”)” to obtain all articles published online in 2013. Titles and abstracts were first scanned for articles presenting original results involving fear conditioning in rodents that were written in English. Selected articles underwent full-text screening for selection of experiments that (a) described the effects of a single intervention on fear conditioning acquisition or consolidation, (b) had a clearly defined control group to which the experimental group is compared to, (c) used freezing behavior as a measure of conditioned fear in a test session and (d) had available data on mean freezing, SD or SEM, as well as on the significance of the comparison. Articles were screened by one of two investigators (C.F.D.C. or T.C.M.) for relevant data and were analyzed by the other – thus, all included experiments were dual-reviewed.

Only experiments analyzing the effect of interventions performed before or up to 6 hours after the training session (i.e. those affecting fear conditioning acquisition or its immediate consolidation) were included. Data on mean freezing and SD or SEM were obtained for each group from the text when available; otherwise, it was extracted using Gsys 2.4.6 software (Hokkaido University Nuclear Reaction Data Centre). When exact sample size for each group was available, the experiment was used for the analysis of effect size and statistical power – otherwise, only effect size was obtained, and the experiment was excluded from power analysis. For individual experiments, study design characteristics were also obtained, including species and sex of the animals, type of conditioning protocol, type, timing and site of intervention.

From each comparison, we also obtained the description term used by the authors in the results session of the paper. Classification of the terms used to describe effects (**S1 and S2 Tables**) was based on a blinded assessment of words or phrases by a pool of 14 researchers who were fluent or native speakers of English and had current or past experience in the field of behavioral neuroscience. Categories were given a score from 0 to 2 in order of magnitude (i.e. 0 = weak, 1 = neutral, 2 = strong for significant results; 0 = similar, 1 = neutral, 2 = trend for non-significant results), and the average results for all researchers was used as a continuous variable for analysis.

Apart from experiment-level variables, we also extracted article-level data such as impact factor of the journal in which it was published (based on the 2013 Journal Citations Report), number of citations (obtained for all articles on August 26^th^ 2016), country of origin (defined by the corresponding author’s affiliation) and the 7 risk of bias indicators described on **Table 1**. For article-level correlations, we compiled these measures into a normalized score.

After completion of data extraction, all calculations and analyses were performed according to the previously specified protocol. Specific details of calculations (as well as the raw data used) are presented as **Supplementary Data**. After this, the following additional analyses were performed in an exploratory fashion:

(a) To confirm that residual freezing levels after memory-impairing interventions were indeed above training values, demonstrating that most amnestic intervention have partial effects, we extracted pre-conditioning freezing levels from training sessions when these were available. These levels were obtained for pre-shock periods only, and separated as baseline values for contextual (i.e. freezing in the absence of tone) or tone conditioning (i.e. freezing in the presence of a tone, but before shock), as displayed in **S5 Fig**. These were compared to the corresponding test session values for treated groups in memory-impairing interventions by an unpaired *t* test based on the extracted means, SD or SEM and sample size when these were available.

(b) In the original protocol, only the mean of all effective interventions (i.e. upper-bound effect size) was planned as a point estimate to be used for power calculations, although we acknowledged this to be optimistic [20]. We later decided to perform power calculations based on the mean effect size of the experiments achieving power above 0.95 on the first analysis (i.e. intermediate effect size) to avoid effect size inflation, as we reached the conclusion that this would provide a more realistic estimate. Additionally, we calculated power based on the mean effect size of the whole sample of experiments as a lower-bound estimate, and presented all three estimates in the results section and figures.

(c) In order to evaluate whether the distribution of effect sizes and statistical power varied if effect sizes were defined as absolute differences in freezing levels instead of relative ones, we repeated the analyses in **Figs. 2, 3** and **4** using absolute differences in **S1 Fig., S6 Fig.** and **S9 Fig.**. This proved to be particularly important to demonstrate that correlations between effect sizes and power were not the consequence of a confounding association of both variables with coefficients of variation. We also repeated power and correlation analyses using effect sizes as standardized mean differences (e.g. Cohen’s d) in **S7 Fig.** and **S10 Fig.**)

(d) To further evaluate the possible impact of the negative correlation between coefficients of variation and freezing levels on our results, we decided to use freezing levels as a covariate in the correlations shown in **Fig. 4**. We also checked whether adding freezing levels as a covariate influenced the statistical analyses in **Fig. 5**, **Fig. 6** and **S11 Fig.**, but as this did not have a significant impact on the results in these figures, we only reported the originally planned analyses.

(e) All of our planned analyses were parametric; after extraction, however, it was clear that some of the data deviated from a normal distribution (especially in the case of power estimates, citation counts and impact factor). Because of this, we performed additional non-parametric analyses for the correlations of citations and impact factor with percentage of significant results, mean normalized effect size, statistical power and study quality score.

(f) In the protocol, we had planned to test correlations between normalized effect sizes and statistical power, mean sample size and absolute freezing levels (using the group with the highest freezing). After analyzing the results, we also decided to correlate normalized effect sizes with coefficients of variation (as this, rather than sample size, seemed to explain the lower power of non-significant results), additional power estimates (as using our original estimate led to a ceiling effect) and different estimates of freezing based on the control group or on the mean freezing of both groups (to compare these forms of normalization with the one we chose).

(g) Due to the correlation of study quality assessment with journal impact factor, we performed an exploratory analysis of the correlation of this metric with each of the individual quality assessment items by performing a Student’s t test (corrected for unequal variances by Welch’s correction) between the impact factors of studies with and without each item.

(h) Because of the additional analyses above, we adjusted the number of comparisons/correlations used as the basis of the Holm-Sidak correction for multiple comparisons. The total numbers used for each correction were 14 for experiment-level comparisons, 17 for article-level comparisons, 28 for experiment-level correlations and 8 for article-level correlations, leading to significance thresholds between 0.003 and 0.05.

